# Principles of Meiotic Chromosome Assembly

**DOI:** 10.1101/442038

**Authors:** Stephanie A. Schalbetter, Geoffrey Fudenberg, Jonathan Baxter, Katherine S. Pollard, Matthew J. Neale

**Author notes:** SAS & GF contributed equally to this work.

## Abstract

During meiotic prophase, chromosomes organise into a series of chromatin loops emanating from a proteinaceous axis, but the mechanisms of assembly remain unclear. Here we elucidate how this elaborate three-dimensional chromosome organisation is underpinned by genomic sequence in *Saccharomyces cerevisiae*. Entering meiosis, strong cohesin-dependent grid-like Hi-C interaction patterns emerge, reminiscent of mammalian interphase organisation, but with distinct regulation. Meiotic patterns agree with simulations of loop extrusion limited by barriers, yet are patterned by convergent transcription rather than binding of the mammalian interphase factor, CTCF, which is absent in *S. cerevisiae*—thereby both challenging and extending current paradigms of local chromosome organisation. While grid-like interactions emerge independently of meiotic chromosome synapsis, synapsis itself generates additional compaction that matures differentially according to telomere proximity and chromosome size. Collectively, our results elucidate fundamental principles of chromosome assembly and demonstrate the essential role of cohesin within this evolutionarily conserved process.

## Introduction and Results

During meiosis, eukaryotic chromosomes are broken, repaired and paired with their homologs followed by two rounds of segregation—a series of events accompanied by dynamic structural changes of the chromosomes. Most prominent is the paired arrangement of pachytene chromosomes into a dense array of chromatin loops emanating from proteinaceous axes linked by a central core, the synaptonemal complex (SC), which is highly conserved across eukaryotes^1,2^. In *S. cerevisiae*, structural components include the meiotic cohesin kleisin subunit, Rec8^3^, the transverse filament, Zip1^4^, the axial/lateral elements, Hop1 and Red1^5,6^, and the pro-DSB factors Rec114-Mei4-Mer2 (RMM)^7,8^. Much of our understanding of meiotic chromosome structure has been deduced from a combination of electron microscopy, immunofluorescence microscopy, and the genome-wide patterns of protein localisation determined by ChIP. However, the link between key meiotic protein complexes, chromosome conformation, and genomic sequence remains uncharacterized.

Chromosome conformation capture (3C) techniques generate maps of pairwise contact frequencies that are snapshots of chromosome organisation. 3C methods were originally applied to assay chromosome conformation in *S. cerevisiae*, including during meiosis^9^. Now they are widely used across a range of organisms and cellular contexts to link 3D organisation directly with genomic sequence^10^, revealing important roles of the Structural Maintenance of Chromosomes (SMCs) cohesin and condensin in genomic organization^11,12^, where they likely mediate chromosome compaction via the process of loop extrusion^13^. Here we return to yeast meiosis to interrogate genome-wide chromosome organisation by Hi-C, elucidate mechanisms of chromosome assembly, and define the role of key meiotic chromosome components, including cohesin and the SC.

Starting with a synchronized G1 population we analysed timepoints encompassing DNA replication, meiotic prophase and both meiotic divisions (Fig. 1a,b,c, Extended Data Fig. 1a,b,c). In G1, we detect strong centromere clustering (Fig. 1a,d) and folding back of the arms at the centromeres (Fig. 1a, **Supplementary Fig. 2**), characteristic of a Rabl conformation^9,14^. During meiosis, centromere clustering is transiently dissolved (3-5h, Fig. 1a,d, Extended Data Fig. 1a); this coincides with a global decrease in inter-chromosomal contact frequency at mid-prophase, reflecting chromosome individualisation. Subtelomeric clustering also decreases during meiotic prophase (Fig. 1a,d, Extended Data Fig. 1a), with no evidence for the transient telomeric bouquet conformation, consistent with prior microscopic analyses^15^.

**Figure 1.**
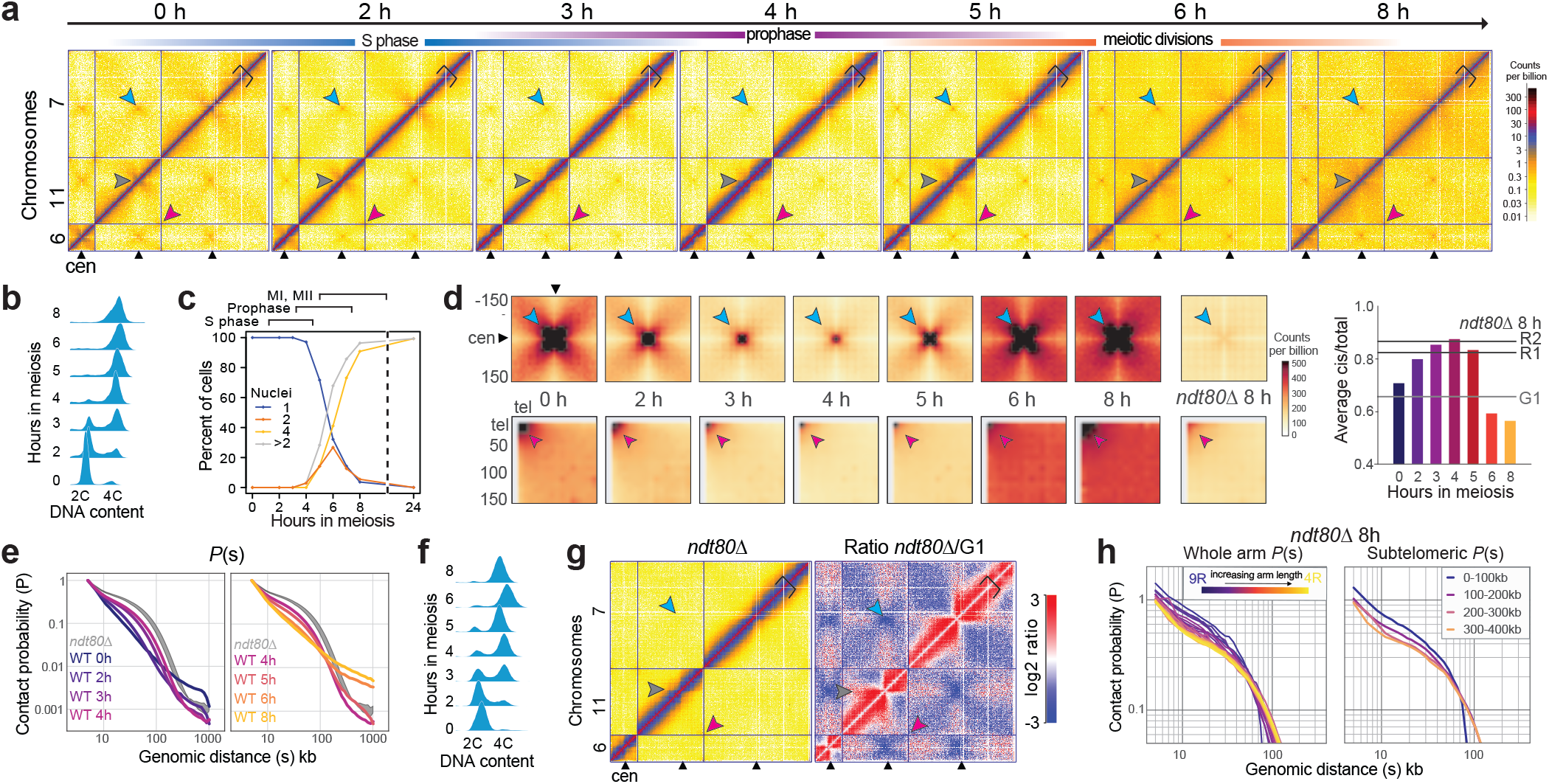
Chromosome conformation during yeast meiosis. **a.** Cells were collected during meiosis at indicated timepoints and analysed by Hi-C. At 0h the cells are in G1. Representative Hi-C contact maps of chromosomes 6, 11, and 7 plotted at 5 kb resolution. Centromeres, telomeres and arm fold-back at the centromere are indicated by blue, red and grey arrows, respectively, and axial compaction by the width of the main diagonal relative to the fixed-width black clamp. For interactive HiGlass^58^ views see: http://higlass.pollard.gladstone.org/app/?config=Z5iwKpjzQpePCXXyvuYGeQ **b.** Meiotic entry assessed by FACS; at 4h, the majority of cells show a 4C peak indicating completion of DNA replication. **c.** Meiotic progression was monitored by quantification of nuclear divisions determined by DAPI staining. Around 4h, cells start to undergo meiotic divisions I and II. The majority of cells undergo meiotic divisions between 4h and 8h, indicating the degree of heterogeneity within the cell population. **d.** *Upper panels*: Average *trans* centromere-centromere contact maps. *Lower panels*: *trans* telomere-telomere contact maps. *Right*: ratio of *cis* to total contact frequency. **e.** Intra-arm contact probability versus genomic distance, *P(s)*, indicating the emergence (*left*) and disappearance (*right*) of chromosome arm compaction during meiosis. Shaded area bounded above and below by the two *ndt80*Δ 8h replicates. **f.** Meiosis was induced in *ndt80*Δ cells for 8h and meiotic entry was checked by monitoring DNA replication by FACS. **g.** *ndt80*Δ cells were grown for 8h in sporulation media and analysed by Hi-C (*left*). Log2 ratio of *ndt80*Δ cells 8h over G1 (*right*). Centromeres and telomeres are indicated by blue and red arrows, respectively, and axial compaction by a black clamp. **h.** *Left*: Contact probability of individual chromosome arms stratified by length. *Right*: Contact probability stratified by the distance from the telomere.

Entering meiosis, contact frequency versus distance, *P(s)*, curves display a shoulder, consistent with the linear compaction of chromosome arms increasing due to *cis*-loop formation (2-4h, Fig. 1e, Extended Data Fig. 1d, e.g. as defined^16^; for review^13^). This change in *P(s)* is reminiscent of the SMC-dependent changes observed via Hi-C during mitosis across species^17–21^. Compaction coincides with meiotic prophase I and the formation of the SC at pachytene, and is lost at later stages (Fig. 1e, Extended Data Fig. 1d).

To study meiotic chromosome conformation in more detail, and to eliminate cell-to-cell heterogeneity (Fig. 1b,c), we enriched for pachytene cells in subsequent experiments by inactivating Ndt80, a transcription factor required for exit from meiotic prophase^22^. *ndt80*Δ cells entered meiosis synchronously, assessed by bulk DNA replication (Fig. 1f), but do not initiate the first nuclear division^22^. Similar to the wild type prophase population (3-5h), but likely accentuated by the increased homogeneity, Hi-C maps of pachytene-enriched cells displayed total loss of centromere clustering (Extended Data Fig. 2) and dramatic chromosome arm compaction (Fig. 1e). Analysing compaction in more detail, shorter chromosomes (Extended Data Fig. 1e) and, in particular, shorter chromosome arms (Fig. 1h, Extended Data Fig. 1f), displayed elevated contact frequency at short genomic separations, and an earlier shoulder, apparently arising from distinct behavior of subtelomeric and subcentromeric regions (Fig. 1h, Extended Data Fig. 1g). Moreover, distinct *P(s)* for chromosomes with different length arms (Extended Data Fig. 1h) suggests that the centromere can insulate the process that leads to differences between arms. In agreement with this, compaction is interrupted at centromeres in Hi-C maps (Fig. 1a, Extended Data Fig. 2b).

**Figure 2.**
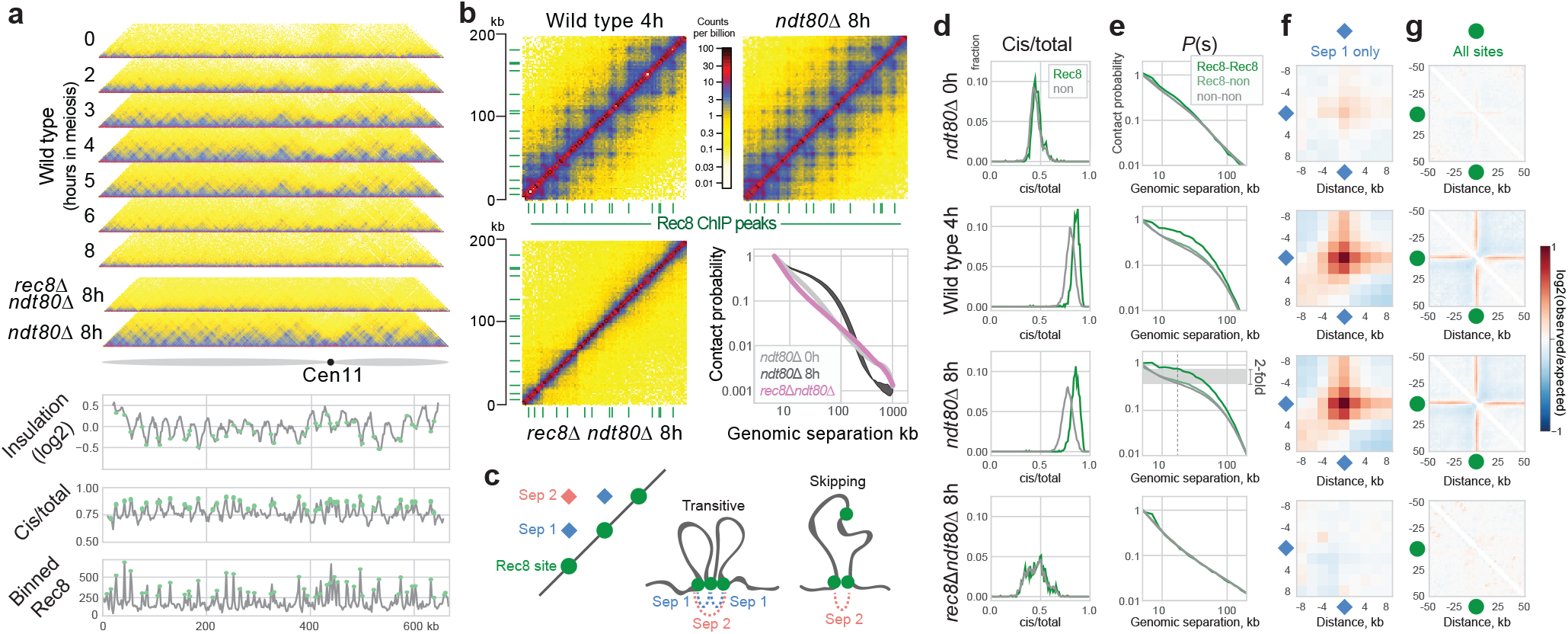
Emergence of a Rec8-dependent grid of punctate interactions in meiosis. **a.** Hi-C contact maps of chromosome 11 for the indicated genotypes plotted at 2 kb bin resolution, showing near-diagonal interactions. Wild type timepoints as in Fig. 1a. **Lower panels**: log2(insulation); cis/total ratio, Rec8 ChIP-seq^27^, all binned at 2kb. Insulation and cis/total calculated from *ndt80*Δ maps. Positions of Rec8 sites indicated as green circles. Genome-wide cis/total (Spearman’s R = 0.62, P < 1e-10) and insulation (R= −0.23, P < 1e-10, insulation window = 20 kb) profiles are correlated with Rec8 occupancy. **b.** Zoom-in into contact maps on chromosome 11 (0-200kb) of wt-4h and *ndt80*Δ (top) and *rec8*Δ (*bottom left*). Contact probability versus genomic distance, *P(s)*, for G1(*ndt80Δ-0h*) and *ndt80*Δ and *rec8*Δ (*bottom right*). Data shown is the average (n = 2) except for wt-4h. Rec8 peak sites called from ChIP-seq data^27^ are indicated in green. For an interactive view see: http://higlass.pollard.gladstone.org/app/?config=Twrh61jGT4SlxotaguTIJg **c.** Simplified illustration of how a grid of peaks on a Hi-C map can emerge between Rec8 sites either by transitive contacts between adjacent loops, or by loops that skip over adjacent sites. Experimentally observed grids extend much further than separation = 2 (Extended Data Fig. 4c) **d.** Cis/total ratios for Rec8 (green) and nonRec8 (grey) sites for indicated datasets. **e.** Contact frequency versus distance between Rec8-Rec8 sites (green), Rec8-nonRec8 sites (light green) and nonRec8-nonRec8 sites (green). **f.** Log2 ratio of contact frequency between adjacent Rec8-sites (separation = 1) compared to average cis interactions. **g.** Log2 ratio of contact frequency centered at Rec8 sites compared to average cis interactions. In *ndt80*Δ, Rec8 sites show: elevated cis/total frequency (0.85 versus 0.77), elevated pairwise contact frequency (∼2-fold at 20 kb), and mild insulation (**f-g**). These distinctions are similar in wild type pachytene (4h) yet absent in G1 (*ndt80Δ-0h*) or in *rec8*Δ.

Zooming in to consider within-arm organization revealed punctate grid-like Hi-C interactions between pairs of loci during prophase (Fig. 2a), particularly prominent in *ndt80*Δ (Fig. 2a,b). Such focal meiotic patterns are more prominent than reported previously^23^—resembling peaks between CTCF sites^24^ rather than topological domains^25,26^ detected in mammalian interphase Hi-C maps— and likely arise from a heterogeneous mixture of ‘transitive’ interactions and ‘skipping’ of peak bases (Fig. 2c).

Genomic regions underlying the punctate Hi-C interactions display a remarkable visual (Fig. 2a,b), and quantitative (Fig. 2d-g), correspondence with previously characterized sites of high Rec8 occupancy^27^. At pachytene, Rec8 sites display elevated cis/total contact frequencies (Fig. 2c), enriched contact frequency (Fig. 2e,f), and evidence of insulation (Fig. 2g)—features that correlate with Rec8 occupancy measured by ChIP (Fig. 2a, **lower**). In wild type cells, Rec8-Rec8 interactions became visible in early prophase (2h), peaked at mid prophase (4h), and were especially prominent in the homogenous *ndt80*Δ cell population (Fig. 2a-g, Extended Data Fig. 4b,c). Importantly, Rec8-Rec8 enrichments are strongest between adjacent sites, decrease between non-adjacent sites with increasing genomic separation, and are absent in *trans* (Extended Data Fig. 4b,c). As for enrichments between CTCF sites in mammalian interphase^28^, these observations argue that a cis-acting process generates such focal interactions in meiosis.

Rec8 is a central component of the meiotic chromosome axis^3^. Assaying a *rec8*Δ mutant enabled us to determine that Rec8 is absolutely required for the emergence of the grid-like Hi-C patterns present in meiosis (Fig. 2a,b). Moreover, *rec8*Δ cells completely lose the shoulder in *P(s)*, indicative of a dramatic loss of arm compaction (Fig. 2b,Extended Data Fig. 4a), similar to that caused by depletion of SMCs in diverse contexts^17,19,21,29–32^. Instead of assembling an axis of loops, *rec8*Δ cells appear to be caught in a state with highly clustered telomeres (Extended Data Fig. 4d, Extended Data Fig. 3), consistent with previous observations by microscopy^33,34^. Moreover, in *rec8*Δ cells *cis* contact frequency is reduced (Fig. 2d), similar to G1 cells, and cis/total no longer correlates with Rec8 occupancy. Instead, *rec8*Δ cis/total displays a decreasing trend along chromosome arms, likely due to persistent telomere clustering (Extended Data Fig. 4d).

**Figure 3.**
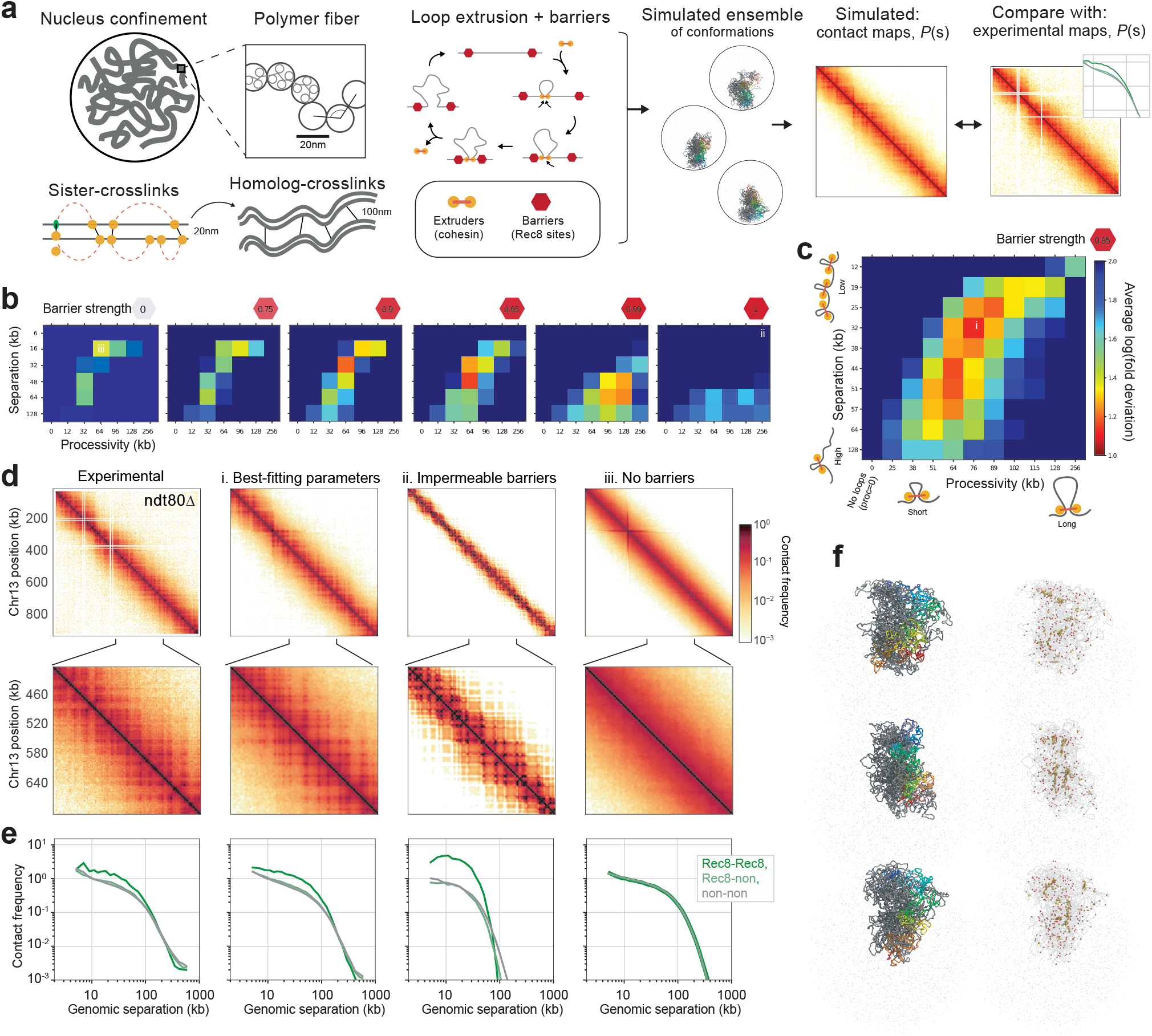
Modelling meiotic chromosome compaction. **a.** In simulations, yeast chr13 was represented as a polymer fiber confined to the nucleus subject to additional meiosis-specific constraints. These include: extruded loops, sister crosslinks, and homolog crosslinks (Methods). Barriers to extruded loops were placed at Rec8 sites^27^. We imposed inter-sister and inter-homologue crosslinks at sites of extruded loop bases in order to approximate the paired arrangement of homologues at pachytene (Extended Data Fig. 6). For each set of extruded loop parameters (*processivity, separation*, and *barrier strength*), conformations were collected and used to generate simulated contact maps. Roughly, *processivity* dictates the size of an extruded loop unimpeded by collisions, *separation* controls the number of active extruders on the chromosome, and *barrier strength* controls the probability that an extruder gets paused when attempting to step past a barrier. Simulated and experimental contact maps were then compared via the combined average fold discrepancy between *P(s)* curves for Rec8-Rec8, Rec8-non, and non-non bin pairs at 2kb resolution. **b.** Goodness-of-fit for indicated barrier strengths over coarse grids of processivity and separation demonstrate that intermediate barrier strengths are required to agree with experimental *ndt80*Δ Hi-C maps. **c.** Goodness-of-fit for a fine grid of processivity versus separation at barrier strength 0.95. Best-fitting models had separation ∼32kb and processivity ∼76kb, corresponding to ∼60% coverage of the genome by extruded loops of average length 26kb. **d.** From *left* to *right*: contact maps for chr13 for *ndt80*Δ, and simulations with (i) best-fitting parameters, (ii) relatively stable loops between neighboring Rec8 sites, and (iii) no barriers. **e.** P(s) split by Rec8-Rec8, Rec8-non, and non-non, as in Fig. 2D. **f.** Conformations for best-fitting simulations, which highlight: (*left*) one chromatid colored from start (red) to end (blue); (*right*) extruders (yellow), extrusion barriers (red), and extruders paused at barriers (orange).

To test how compaction and grid-like interaction patterns could jointly emerge in meiosis, we developed polymer simulations (Fig. 3a,Methods) similar to those used to successfully describe the assembly of TADs in mammalian interphase chromosomes^13^. Importantly, these simulations employ the *cis*-acting process of loop extrusion, where extruders form progressively larger chromatin loops, unless impeded by adjacent extruders or barrier elements (Fig. 3a). Extrusion dynamics are controlled by parameters dictating the *processivity* (average loop size) and *separation* (number of active extruders), as well as the *strength* of barriers (Methods). Because the accumulation of Rec8 at ChIP-seq sites^27^ concomitant with convergent transcription^35^ is indicative of barriers to extrusion^28^, we positioned bi-directional barriers at Rec8 sites.

Simulations were used to explore variations in loop extrusion dynamics to determine whether specific parameter combinations are able to generate Hi-C maps that agree with experimental observation (Fig. 3,Methods). Models with excellent fits were identified in which ∼65% of the genome is covered by extruded loops (Fig. 3b,c, Extended Data Fig. 5)—a far denser array than present in *S. cerevisiae* mitosis^21^, but still less compact than human mitotic cells^20^. Even though extrusion can generate compaction independently of barriers (Fig. 3d), an intermediate barrier strength is essential to match the grid-like patterns observed experimentally (Fig. 3b). Despite the simplifying assumptions, simulated chromosomes displayed many features observed experimentally: (i) chromosomes fold into a loose polymer brush^3,36,37^, with a Rec8-rich core^3^ (Fig. 3f, Extended Data Fig. 5 a); (ii) a grid-like interaction pattern naturally emerges in simulated Hi-C maps (Fig. 3d); (iii) importantly, because loop extrusion is a *cis*-acting process, pairs of Rec8 sites at increasing separations naturally have lower contact frequency (Fig. 3e).

Simulations also highlight the stochasticity of loop positions across the cell population, with most barriers (73%) unoccupied by an extruder, and extruders paused with barrier elements on both sides only a minority of the time (15%) in the best fitting models (Extended Data Fig. 6 c). Because of this, the majority (65%) of extruded loops cross over Rec8 sites, consistent with an average loop size roughly twice the average distance between Rec8 ChIP peaks (26 kb versus 12 kb, Extended Data Fig. 6 d), and remarkably consistent with estimates made using EM (∼20kb^36^). Most strikingly—despite the prominence of Rec8-dependent grid-like features in the experimental data (Fig. 2c)—our simulations indicate that Rec8 sites are not always occupied by extruding cohesins and thus are present at the meiotic chromosome core in only a subset of cells, as inferred previously^38^.

The range of loop extrusion parameters we explored encompasses the situation where Rec8 sites always halt extrusion and *cis*-loops are formed between each consecutive Rec8 site. However, simulations with these parameters have quantitatively poor fits with experimental maps (Fig. 3d e, ii): the bend in *P(s)* comes too early to recapitulate experimental *P(s)*, and Rec8-Rec8 contacts are much too strong. The poor fit of such ‘direct-bridging’ simulations underscores the conclusion that only a fraction of Rec8 sites are occupied in a given cell, and argues that cohesin-dependent *cis*-loops must link regions that are not primary Rec8 binding sites in order to provide compaction without making Rec8-Rec8 enrichments overly strong.

A crucial prediction of our loop extrusion simulations is that depletion of extruders in meiosis would lead to both decompaction and loss of the grid-like pattern of Hi-C interactions. When we repeated our fitting procedure for *rec8*Δ, the best fits were for simulations with either no, or very few, extruded loops (Extended Data Fig. 5e). The lack of compaction in these simulations (Extended Data Fig. 5a) is consistent with previous EM showing decompacted chromatids in *rec8*Δ^3^. Such joint consistency between Hi-C and imaging data further supports loop extrusion as a mechanism underlying assembly of the cohesin-rich core and contributing to chromosomal compaction in meiosis. Our simulations also open the possibility that overly-shortened axes observed upon Wapl^39,40^ and Pds5^41^ depletion may reflect heightened extruder processivity^42^ upon which shortened SCs are assembled, and predict that such perturbations would cause a rightward shift in the *P(s)* shoulder measured via Hi-C (Extended Data Fig. 5c).

To investigate how homologue synapsis affects chromosome conformation, we assayed pachytene cells in the absence of Zip1, the transverse filament of the SC^4^, and Hop1, an axial element required for Zip1 loading^6^ (Fig. 4a,b). Both *zip1*Δ and *hop1*Δ Hi-C maps retained the Rec8-dependent punctate interactions (Fig. 4b, Extended Data Fig. 2b,c), and displayed compaction relative to G1 or *rec8*Δ, but with the *P(s)* shoulder shifted left relative to *ndt80*Δ (Fig. 4c). Attempts to model the known *zip1*Δ and *hop1*Δ defects in chromosome synapsis simply by removing interhomologue crosslinks from best-fitting *ndt80*Δ simulations did not recapitulate the *P(s)* shift observed experimentally (Extended Data Fig. 5 f), consistent with the suggestion that interhomologue contacts make only a minor contribution within meiotic Hi-C maps^23^. Instead, best-fitting simulations had shifts towards slightly lower processivity and larger separation, consistent with less axial compaction relative to the *ndt80*Δ control (Fig. 4e). Interestingly, subtelomeric regions no longer displayed a distinct *P(s)* in *zip1*Δ and *hop1*Δ (Fig. 4d), suggesting that chromosome compaction at chromosome termini is regulated differentially.

**Figure 4.**
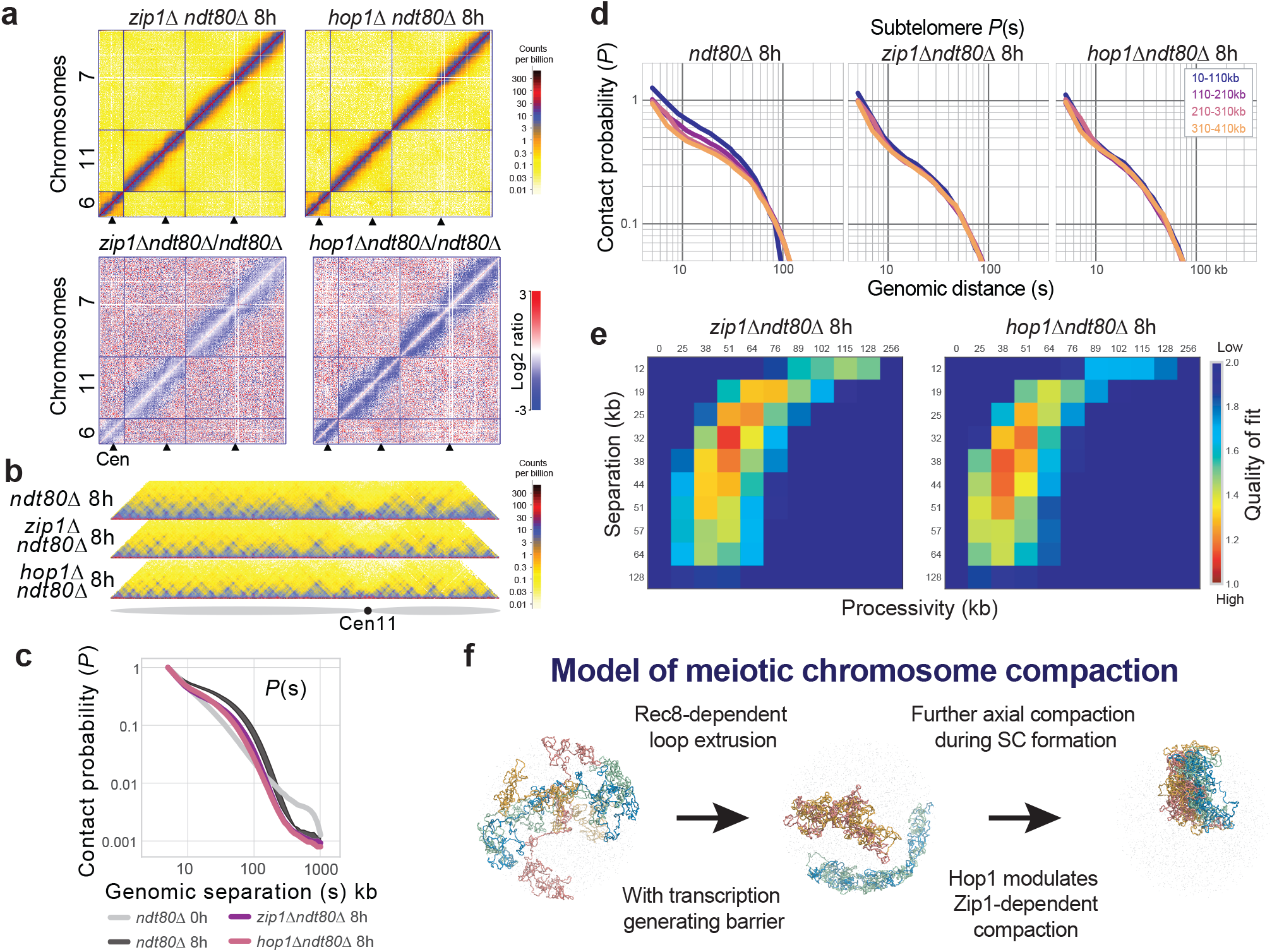
Hop1 and Zip1-dependent compaction of Rec8-dependent loops. **a.** *Top*: Hi-C maps for *hop1*Δ and *zip1*Δ (plotted as in Fig. 1a). *Bottom*: Log2 ratio of *hop1*Δ over *ndt80*Δ (as in Fig. 1g). For interactive views of the full genome, see http://higlass.pollard.gladstone.org/app/?config=TTBGu5DDR0SHAa09zrjTXA **b.** Hi-C contact maps of chromosome 11 for *hop1*Δ and *zip1*Δ plotted at 2kb bin resolution, showing near-diagonal interactions, as in Fig. 2a. **c.** Contact probability versus genomic distance for G1, *ndt80*Δ, *hop1*Δ, *zip1*Δ. Shaded area bounded above and below by *ndt80*Δ replicas. Average between two replicas for *zip1*Δ and one sample for G1 and *hop1*Δ are shown. **d.** Contact probability over genomic distance averaged over all chromosome arms stratified by distance from the telomere. **e.** Goodness-of-fit for simulations without homolog crosslinks with a fine grid of processivity versus separation at barrier strength 0.95 *zip1*Δ and *hop1*Δ. **f.** Model of meiotic chromosome compaction: Rec8-dependent loop formation leads to initial chromosome arm compaction and emergence of a grid-like pattern of Hi-C interactions that jointly agrees with a mechanism of loop extrusion including barrier elements. We suggest that transcription could impose such barriers. Hop1 and Zip1 are dispensable for this step, but are required for synapsis, where additional compaction occurs differentially along chromosome arms.

## Discussion

Our analysis of meiotic chromosome organisation via Hi-C reconciles the function and localisation of factors thought to shape meiotic chromosomes with their 3D organisation, revealing the emergence of a punctate grid of interactions concomitant with initial stages of meiotic chromosome compaction. Crucially, we formally demonstrate the link between preferential positioning of meiotic cohesin along the genome^27,35^ and the inference that these loci come into close proximity based on the localization of Rec8 to the chromosomal axes^3^. Remarkably, the punctate cohesin-dependent interactions in yeast meiosis emerge despite the absence of CTCF in this organism; this challenges previous models where focal Hi-C peaks are strictly dependent on CTCF^24,31,43^, and indicates that alternative mechanisms of loop positioning must exist.

Notably, whilst much less prominent, locus-specific folding is evident in equivalent high-resolution Hi-C maps of mitotic cells (Extended Data Fig. 7)—something that was hidden within lower-resolution analyses^19,21^. The correspondence between cohesin positioning and convergent transcription in both meiosis^35^ and mitosis^44,45^ argues that transcription may be a fundamental and ubiquitous process capable of broadly patterning locus-specific chromosome organisation by modulation of cohesin dynamics^46^. Indeed, the stronger meiotic patterns are particularly reminiscent of the extended grid-like Hi-C patterns observed in interphase mammalian cells upon depletion of the cohesion unloader, Wapl^31,47^, wherein “vermicelli”-like chromatids arise with a cohesin-rich backbone^48^, emphasising the influence of cohesin dynamics on loop extrusion. We favour the view that transcription acts as a barrier to cohesin-dependent loop extrusion, rather than as a motive force as previously proposed^35,49,50^, consistent with transcription-independent compaction by cohesin in mammalian interphase^47^ and direct observation of extrusion by the related SMC condensin *in vitro*^51^.

Consistent with the idea that large DNA-protein complexes, like a kinetochore, can impede extrusion, we observe a paucity of Rec8-dependent loops spanning the centromere. Nevertheless, whether barriers arise directly from Pol2 binding, or indirectly via other axis proteins, remains to be determined. Indeed, the reason for the why loops are more strictly positioned in meiosis compared to mitosis is intriguing. However, our observations enable us to rule out the axial element, Hop1, the SC lateral element, Zip1, and the process of homologous recombination mediated by Spo11, Sae2, and Dmc1 (unpub. obs.) as important for the generation of such patterns.

Our simulations also reveal a nuanced picture of meiotic chromosome assembly: loops are, on average, larger than the inter-Rec8 peak distance, and more than half of the loop bases are not associated with preferred sites of Rec8 binding. Moreover, it is likely that loop sizes and positions vary widely from one cell to another, making classifications of genomic regions as ‘axis’ or ‘loop’ a great oversimplification. The agreement between our simulations and experimental data furthers the case for loop extrusion as a general mechanism^19–21,23,28,52–55^ that is flexibly employed and regulated in interphase, mitosis, and meiosis.

Our results also reveal how the interplay between the synapsis components, Hop1 and Zip1, influences chromosome morphology. That Hop1 and Zip1 are both required to increase chromosome compaction at pachytene likely points at their joint role in promoting synapsis^4,6^, and supports the view that synapsis itself modulates axial compaction. Interestingly, whilst Zip1 binds largely uniformly along the arms of pachytene chromosomes^56^, subtelomeres and short chromosomes display an increase in short-range contacts and an earlier shoulder in *P(s)*, consistent with smaller loops or less compression of spacers between loops in these regions, and therefore less axial compaction. Because such differences correlate with disproportionate retention of Hop1 in these regions^56^ and diminished efficiency of synapsis^57^, it is possible that Hop1 impedes the pathway whereby Zip1 imposes additional compaction upon synapsis. Nevertheless, it is unclear whether Zip1 mediates this effect by modifying loop extrusion dynamics, or via a distinct process of axial compression, as has been argued for higher eukaryote mitotic chromosome compaction^20^. Given the influence that chromosome structure has over so many aspects of meiosis, teasing apart these mechanisms is of great future interest.

## Acknowledgements & Contributions

SAS and MJN planned the study, performed wet-lab work and data analysis. GF and KSP developed polymer simulations and performed data analysis. SAS, GF, JB, KSP and MJN discussed results and wrote the manuscript. We thank Tim Cooper for deploying the HiC Pro installation, Scott Keeney and Franz Klein for sharing S. cerevisiae strains, Svetlana Lyalina for assistance with QB3 GPUs, Anton Goloborodko for suggesting the use of looplib, and Nezar Abdennur for feedback.

## Data & Code Availability

All processed Hi-C matrices are publicly accessible via the interactive HiGlass viewer58. Live web links are provided with each figure. Raw sequence reads will be made publicly accessible via the SRA repository upon final publication. Code for data analysis is either already publicly available online or will be made available on request or via Github upon final publication.

## Funding Statement

SS and MJN are supported by an ERC Consolidator Grant (#311336), the BBSRC (#BB/M010279/1) and the Wellcome Trust (#200843/Z/16/Z). KP and GF are supported by NIMH grant #MH109907, NHLBI grant #HL098179, and the San Simeon Fund.

## Methods

### Yeast strains and cell culture growth

Strains used in this study were derived from SK1 and are listed in Table 1.

**Table 1.**
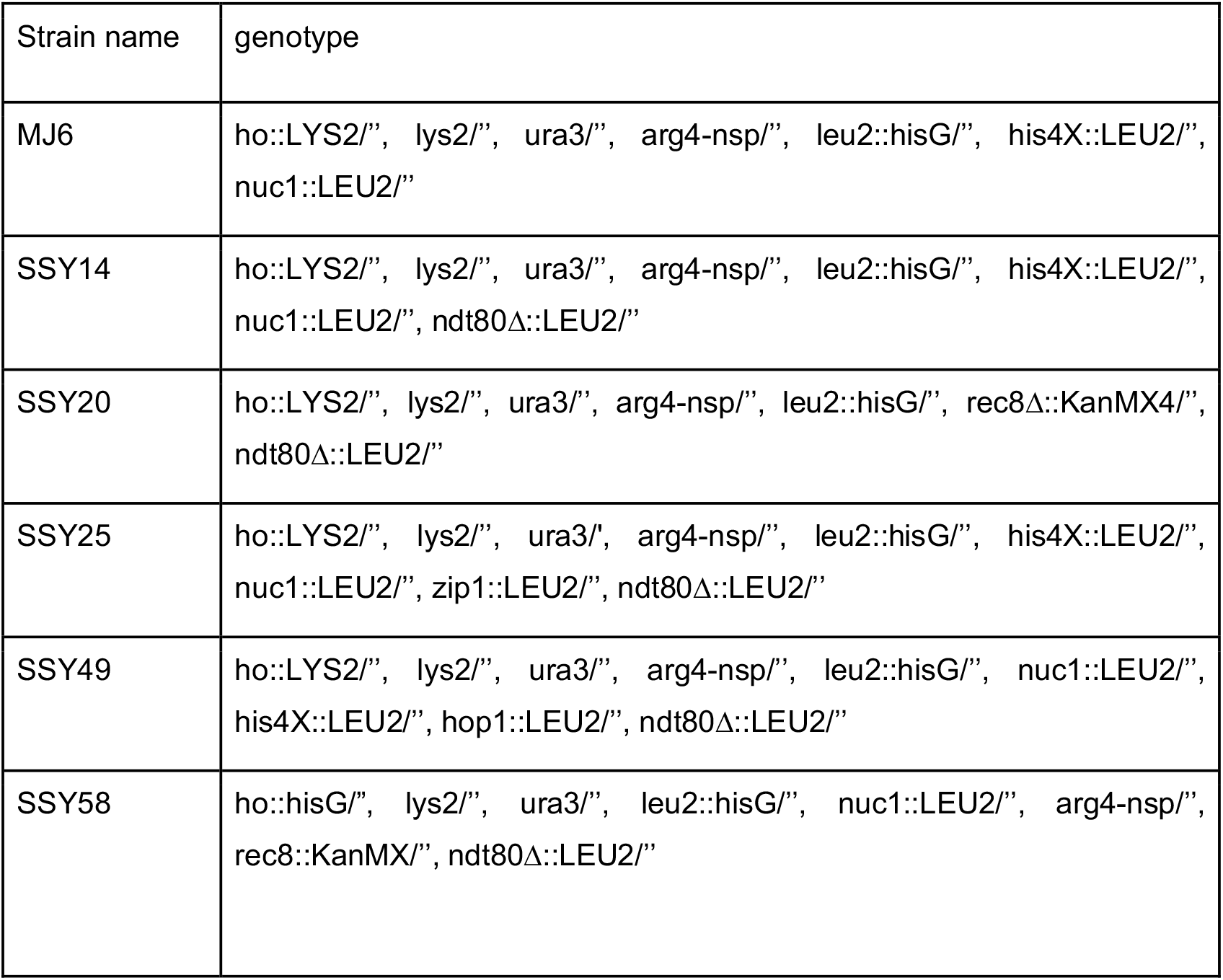
S. cerevisiae strains used in this study

### Monitoring meiotic progression by flow cytometry and quantification of nuclear divisions

Cells were fixed in 70% EtOH, digested with 1 mg/ml RNAse (10 mM Tris-HCl pH 8.0, 15 mM NaCl, 10 mM EDTA pH 8.0) for 2 h at 37 °C, 800 rpm and subsequently treated with 1 mg/ml Proteinase K in 50 mM Tris-HCl pH 8.0 at 50 °C, 800 rpm for 30 min for analysis by FACS. Cells were then washed in 50 mM Tris-HCl pH 8.0 and stained in the same buffer with 1 uM Sytox green overnight in the fridge. FACS profiles were plotted with R using the library *hwglabr2* (https://github.com/hochwagenlab/hwglabr2). Fixed cells were also used for quantification of nuclear divisions by spreading onto a microscope slide, mounting with Fluoroshield containing DAPI followed by analysis with a Zeiss Scope.A1 microscope.

### Hi-C library preparation

The Hi-C protocol used was amended from59 by ∼5-fold reduction in all materials and volumes. Briefly, S. cerevisiae diploid cells were synchronised in G1 by growth at 30 °C for ∼15 h in 30 ml YPA (1% Yeast extract, 2% Peptone, 1% K-acetate) to OD600 of ∼4, harvested, washed, and resuspended in prewarmed sporulation medium (2% KAc with 0.2x nutritional supplements) before fixing 5 ml aliquots (20-30 ODs) of relevant timepoints with formaldehyde at 3% final concentration for 20 min at 30 °C, 250 rpm, then quenched by incubating with a final concentration of 0.35 M Glycine (2x the volume of Formaldehyde added) for an additional 5 minutes. Cells were washed with water split into two samples and stored at −80°C ready for library preparation. Cells were thawed, washed in spheroplasting buffer (SB, 1 M Sorbitol, 50 mM Tris pH 7.5) and digested with 100 ug/ml 100T Zymolyase in SB containing 1% beta-Mercaptoethanol for 15-20 min at 35°C. Cells were washed in restriction enzyme buffer, chromatin was solubilised by adding SDS to 0.1% and incubating at 65 °C for 10 minutes. Excess SDS was quenched by addition of Triton X100 to 1%, and chromatin was incubated with 2.07U/ul of *Dp*nII overnight at 37 °C. DNA ends were filled in with nucleotides, substituting dCTP for biotin-14-dCTP using Klenow fragment DNA polymerase I at 37 °C for 2 h followed by addition of SDS to 1.5% and incubation at 65 °C for 20 min to inactivate Klenow and further solubilise the chromatin. The sample volume was diluted 15-fold, crosslinked DNA ends ligated at 16 °C for 8 h using 0.024U/ul of T4 DNA ligase, and crosslinks reversed by overnight incubation at 65 °C in the presence of proteinase K. DNA was precipitated with ethanol, dissolved in TE and passed through an Amicon 30 kDa column. DNA was further purified by phenol:chloroform:isoamylalcohol extraction and precipitated again before treating with RNaseA at 37 °C for 1 h. Biotin was removed from unligated ends by incubation with T4 DNA polymerase at 20°C for 4 h and at 75 °C for 20 min for inactivation of the enzyme. DNA was subsequently fragmented using a Covaris M220 (Duty factor 20%, 200 cycles/burst, 350s, 20 °C), and DNA ends were repaired and A-tailed using T4 DNA polymerase, T4 Polynucleotide Kinase and Klenow fragment DNA polymerase I before isolating fragments of 100-250 bp using a Blue Pippin (Sage). Biotinylated fragments were enriched using streptavidin magnetic beads (C1) and NextFlex (Bioo Scientific) barcoded adapters were ligated while the DNA was on the beads. Resulting libraries were minimally amplified by PCR and sequenced using paired end 42 bp reads on a NextSeq500 (Ilumina; Brighton Genomics).

### Hi-C data processing and analysis

Hi-C sparse matrices were generated at varying spatial resolutions using the Hi-C-pro pipeline^60^, using a customised S288c reference genome (*SK1Mod*, in which high confidence SK1-specific polymorphisms were inserted in order to improve read alignment rates; manuscript in preparation) and plotted using R Studio (version 1.0.44) after correcting for read depth differences between samples. Raw read statistics are presented in Table 2. Repeat biological samples gave broadly similar matrices and, unless indicated otherwise, were averaged to improve their expected quantitative accuracy. As visual inspection indicated a number of potential translocations in the SK1 strain as compared with the S288c reference genome, for conservative downstream analyses, additional bins were masked if they contained potential translocations. Such bins were identified if they either had values in *trans* at the level of the median of the third diagonal in *cis*, or the maximum value in *trans* exceeded the maximum value in *cis* for SSY14 for bins displaying these properties in either *ndt80D-0h* or in *ndt80D-8h* and for MJ6 in *wt-0h* or *wt-4h*. chr1 was excluded from downstream analysis as few informative bins remained after filtering potential translocations.

**Table 2.** Hi-C Libraries.

| **Name** | **mutations** | **Sample name** | **valid pairs (M)** |
| --- | --- | --- | --- |
| **Main figures** |  |  |  |
| wt-0h/G1 |  | HiC_MJ6_wt_2A_0h | 14.5 |
| wt-2h |  | HiC_MJ6_wt_2A1_2h | 27.6 |
| wt-3h |  | HiC_MJ6_wt_2A_3h | 24.1 |
| wt-4h |  | HiC_MJ6_wt_2A_4h | 28 |
| wt-5h |  | HiC_MJ6_wt_2A1_5h | 27.6 |
| wt-6h |  | HiC_MJ6_wt_2A1_6h | 27.6 |
| wt-8h |  | HiC_MJ6_wt_2A3_8h | 19 |
| *rec8*Δ | *rec8*Δ *ndt80*Δ | average |  |
| *rec8*Δ replica 1 | *rec8*Δ *ndt80*Δ | HiC_SSY20_ndt80Drec8D_1A2_8h | 39.3 |
| *rec8*Δ replica 2 | *rec8*Δ *ndt80*Δ | HiC_SSY58_ndt80Drec8D_2A_8h | 20.2 |
| *ndt80*Δ | *ndt80*Δ | average, 8h |  |
| G1 | *ndt80*Δ | HiC_SSY14_ndt80D_1A2_0h | 36 |
| *ndt80*Δ-4h | *ndt80*Δ | HiC_SSY14_ndt80D_1A_4h | 11.9 |
| *ndt80*Δ replica 1 | *ndt80*Δ | HiC_SSY14_ndt80D_1A1_8h | 22.9 |
| *ndt80*Δ replica 2 | *ndt80*Δ | HiC_SSY14_ndt80D_2A2_8h | 37 |
| *zip1*Δ | *zip1*Δ *ndt80*Δ | average |  |
| *zip1*Δ replica 1 | *zip1*Δ *ndt80*Δ | HiC_SSY25_ndt80Dzip1D_1B2_8h | 22.7 |
| *zip1*Δ replica 2 | *zip1*Δ *ndt80*Δ | HiC_SSY25_ndt80Dzip1D_2A_8h | 28.6 |
| *hop1Δ* | *hop1Δ ndt80Δ* | hop1 ndt80 |  |
| *hop1Δ* replica1 | *hop1Δ ndt80Δ* | HiC_SSY49_ndt80Dhop1D_1A_8h | 32.8 |
| **Supplementary figures** |  |  |  |
| wt-2h |  | HiC_MJ6_wt_3A_2h | 22.5 |
| wt-3h |  | HiC_MJ6_wt_3A_3h | 19.8 |
| wt-4h |  | HiC_MJ6_wt_3A_4h | 16.7 |
| wt-6h |  | HiC_MJ6_wt_3A_6h | 37.6 |

**Table 3.**
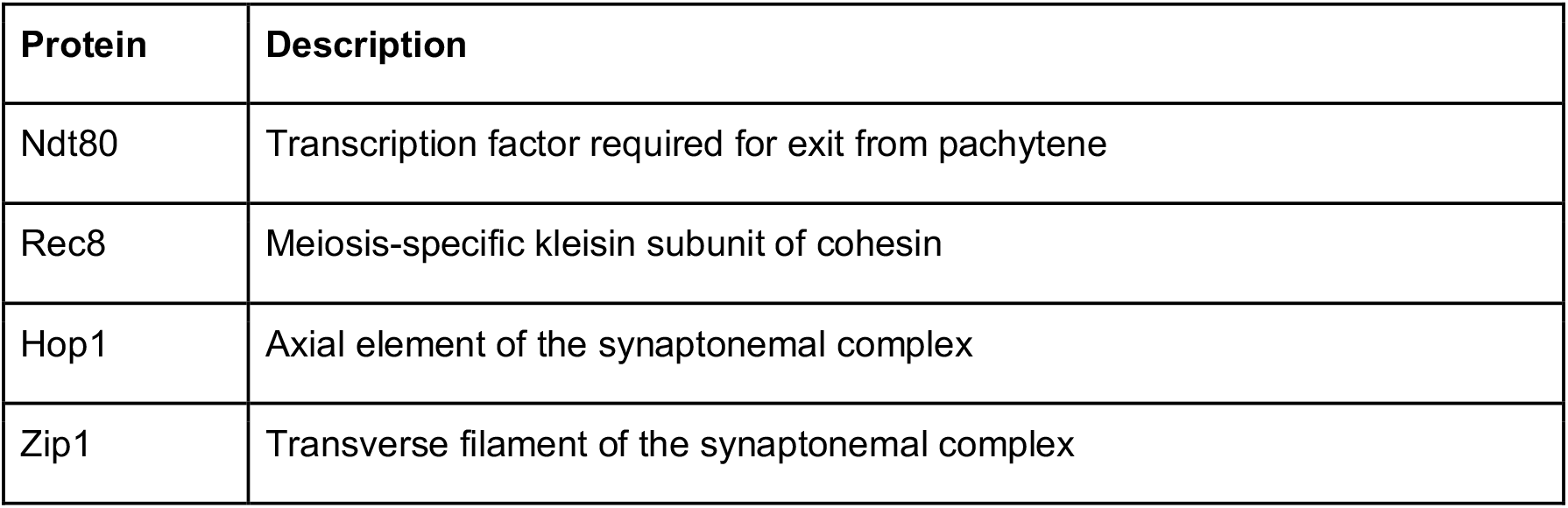
Overview of proteins described in this study.

Average maps centered at centromeres and telomeres were calculated as in^61^, ensuring that collected patches for average centromere maps did not extend inter-chromosomally, and collected patches for average telomere maps did not extend beyond centromeres or inter-chromosomally. Contact frequency versus distance curves, *P(s)*, were calculated from 2 kb binned maps, with logarithmically-spaced bins in *s* (numutils.logbins, https://bitbucket.org/mirnylab/mirnylib, start = 2, end = max(binned arm lengths), N = 50), and restricting the calculation to bin pairs within chromosomal arms and excluding bins less than 20 kb from centromeres or telomere (as in^61^), and normalized to the average value at 4 kb. *P(s)* stratified by distance to telomeres was calculated using the combined distance to telomeres for each bin-pair (as in^62^), and excluded bins-pairs where one bin was closer to a centromere than telomere along that arm. Distance to centromeres, and *P(s)* stratified by this distance, was calculated similarly. Log2 insulation profiles were calculated using a sliding diamond window (as in^63^) with a +/-20 kb (+/-10 bins) extent; as in^43^ downstream analyses were restricted to when there were zero or one filtered bins in the sliding window. To calculate histograms of cis/total (Fig. 2D), bins were defined as either Rec8 or non-Rec8. To calculate *P(s)* split by Rec8 bin-pair status, each bin-pair (i.e. entry of the heatmap) was assigned as either Rec8-Rec8, Rec8-nonRec8, or non-non (e.g. Fig. 2E). *P(s)* was then aggregated separately across chromosomes for these three categories, similar to calculation of *P(s)* within and between TADs^28^. Average log2 observed/expected maps were calculated by first dividing by intra-arm *P(s)* and then averaging together appropriate patches of Hi-C maps. Correlations between Rec8 occupancy from^27^ and insulation or cis/total profiles excluded chromosome 12 because the rDNA locus greatly alters the insulation profile within the right arm of the chromosome.

### Polymer simulations

Meiotic loop extrusion simulations begin with a generic polymer representation of the yeast chromatin fiber similar to that used in previous models of yeast mitotic chromosomes^21^, where each 20 nm monomer represents 640 bp (∼4 nucleosomes). We simulated the chromatin fiber with excluded volume interactions and without topological constraints, using Langevin dynamics in OpenMM, as in^64,65^. Importantly, meiotic simulations remove the geometric constraints specific to the Rabl conformation^66,67^ because this is not visible in meiotic pachytene *ndt80*Δ Hi-C maps. As our focus was to characterize the grids of intra-chromosomal interactions, we considered a system with multiple copies of chromosome 13, equivalent to four copies of the haploid genome in terms of total genomic content (4 × 13 copies of chromosome 13), to enable efficient computational averaging of simulated Hi-C maps. Extruded loops were generated according to parameters that describe the dynamics of loop extruders, using the simulation engine described in^68^: extruder *separation*, extruder *processivity*, chromatin fiber *relaxation time* relative to extruder velocity, and *barrier strength*. Because yeast chromosomes are short compared to higher eukaryote chromosomes, relaxation time is relatively rapid and we focused on *separation, processivity*, and *barrier strength*. At every given timepoint an extruded loop is realized as a bond between monomers at the two bases of the loop (see ./src/examples/loopExtrusion in https://bitbucket.org/mirnylab/openmm-polymer/).

Upon encountering a barrier, a loop extruder is paused with probability according to the barrier strength; barrier strength = 1 indicates an impermeable barrier, barrier strength = 0 indicates no impediment to extrusion. We assume loop extrusion occurs independently on each chromatid, and simulate loop extrusion dynamics on a 1D lattice (as in^28^) where the number of lattice sites equals the total number of monomers (75,140). Bi-directional barriers were placed at monomers with positions corresponding to Rec8 ChIP-seq sites^27^, and pause extruders according the barrier strength parameter. We assume a uniform birth probability, constant death probability, and that all barriers have an equal strength; as additional data becomes available, these assumptions can be relaxed and more detailed models can be built.

We investigated scenarios where chromatids are then either left individualized (52 copies), crosslinked to sisters (26 pairs), or additionally paired with homologs (13 pairs-of-pairs). For simulations with sister crosslinks, these were added (following^54^) when extruded loop bases were present at cognate positions ± 30 monomers (∼20kb) on both chromatids (distance = 20nm); homolog crosslinks were added similarly when sister crosslinks were present on both chromatids (distance = 100nm); centromeres and telomeres were always paired, and both presented impermeable (strength = 1) boundaries to extruders. To avoid introducing pseudo-knots, if extruded loops were nested only the outer cohesins were considered as possible bases for sister crosslinks, sister crosslinks were only allowed between the same side of loop bases (i.e. left-to-left arm or right-to-right arm), and sister crosslinks were only added between bases at the reciprocal minimum distance.

For calculation of simulated Hi-C maps, contacts were recorded from conformations of the full system, which includes intra- and inter-sister, and interhomologue contacts. Because experimental Hi-C here does not distinguish either sisters or homologs, contacts were then aggregated into one simulated map. For each model and parameter set we investigated, we collected an ensemble of conformations, generated simulated chr13 Hi-C maps, and compared their features and *P(s)* with those from experimental Hi-C maps. Each simulated chr13 map represented an average over 5400 conformations. *P(s)* for chr13 was calculated from 2kb binned simulated maps exactly as for experimental maps. Maps of goodness-of-fit between simulations and experimental data (e.g. Fig. 3b,c) were computed as the geometric standard deviation of the ratio of simulated to experimental *P(s)* combined across *P^Rec8-Rec8^(s)*, *P^Rec8-non^(s)*, and *P^non-non^(s)*, as was previously done for P(s) within TADs of multiple sizes and between TADs^28^, for *s* from 10kb to 300kb. This measure reflects the typical fold-deviation for *P(s)*.

Simulated ChIP-seq profiles (Extended Data Fig. 6b) for Rec8 were generated by aggregating the position of extruded loop bases (two per extruded loop) across conformations. Statistics of extruded loop positioning relative to Rec8 sites was calculated with *loopstats.py* in *looplib* (https://github.com/golobor/looplib), and arc diagrams (Extended Data Fig. 6c) with *loopviz.py*. Conformations showing chromatids or positions of extruded loop bases were rendered in PyMOL (https://pymol.org/sites/default/files/pymol.bib).

---

**Extended Data Fig. 1.**
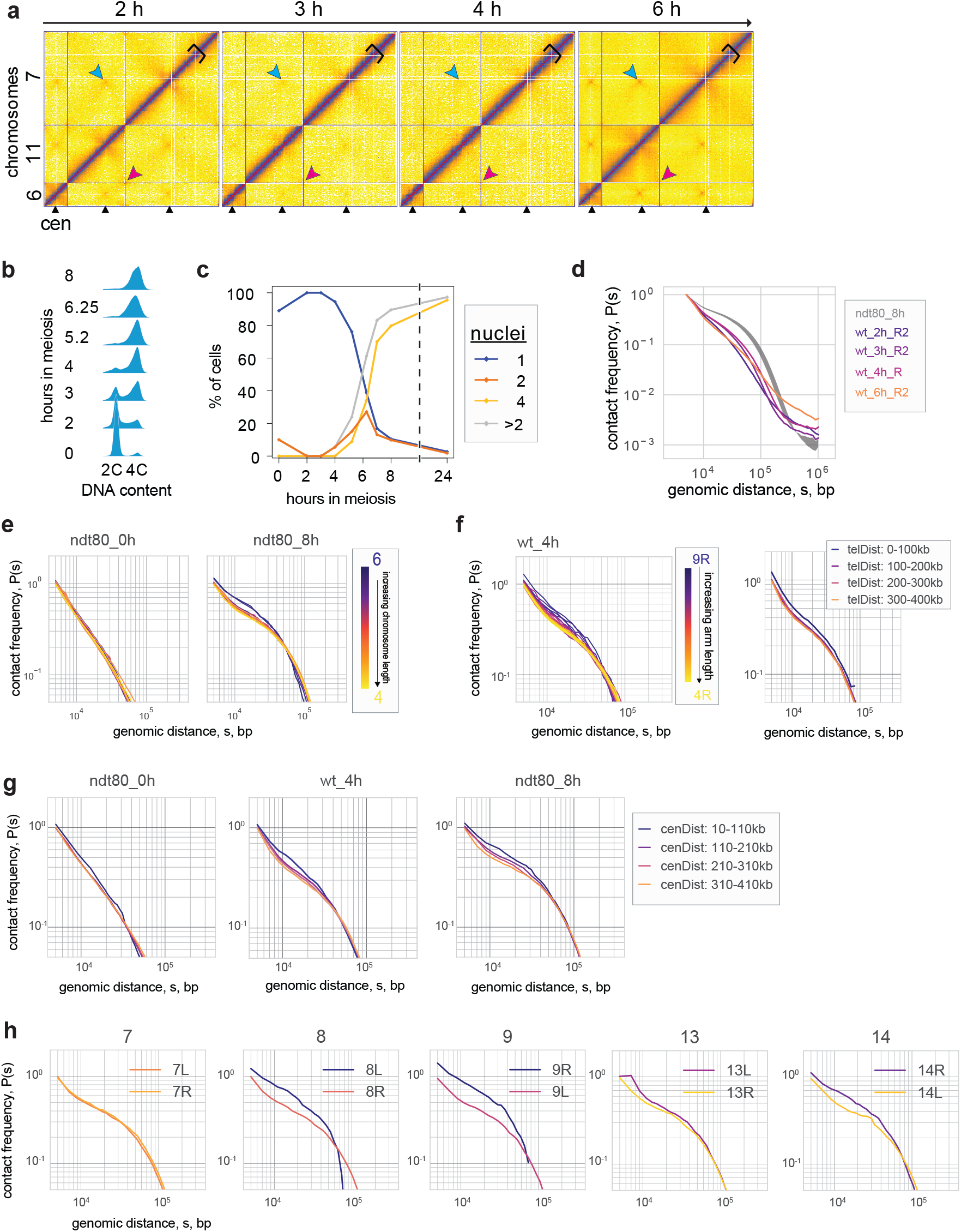
Temporal and chromosome length-specific analysis of meiotic chromatin conformation. **a-d.** Results from a replicate timecourse, collected and characterized independently of the timecourse in Fig. 1. **a.** Hi-C maps, plotted as in Fig. 1a. **b.** FACS as in Fig. 1b. **c.** DAPI as in Fig. 1c. **d.** *P(s)* as in Fig. 1e. **e.** *P(s)* for chromosomes stratified by size for *ndt80*Δ-0h, *ndt80*Δ-8h. Short chromosomes display relatively elevated *P(s)* at short distances, and an earlier shoulder. **f.** *Left*: *P(s)* for individual chromosome arms, stratified by size for wt-4h. Short arms display relatively elevated *P(s)* at short distances, and an earlier roll-over. *Right*: Intra-arm *P(s)* stratified by the distance from the telomere for wt-4h, averaged across all chromosomes. Telomere-proximal regions display elevated *P(s)* at short distances. **g.** Intra-arm *P(s)* stratified by the distance from the centromere for G1 (*ndt80*Δ-0h), *wt-*4h, *ndt80*Δ-8h, averaged across all chromosomes. **h.** Contact probability of single chromosome arms for *ndt80*Δ-8h.

**Extended Data Fig. 2.**
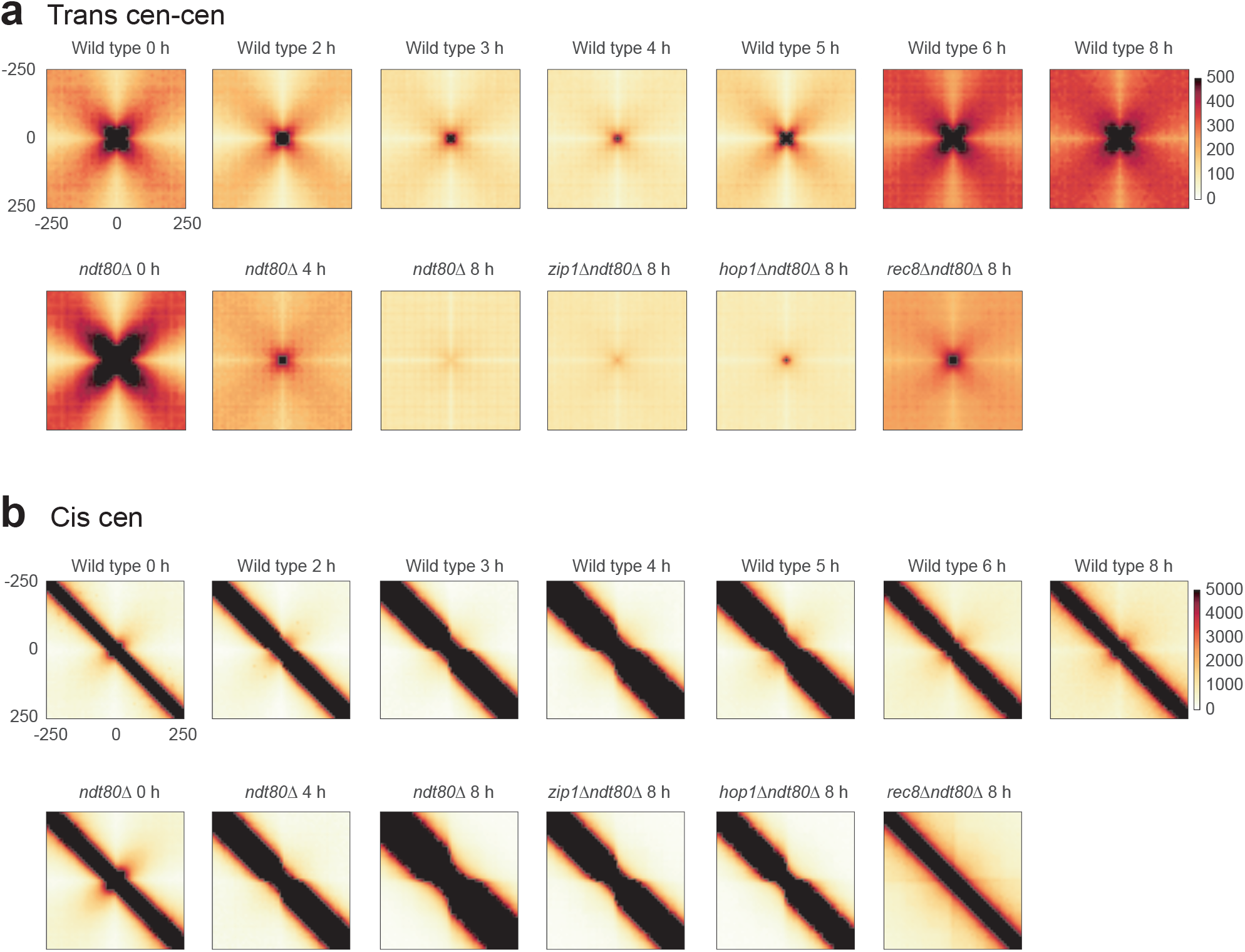
Aggregate analysis of centromeric interactions in meiosis. **a.** Average *trans* centromere-centromere contact maps for indicated data sets. **b.** Average *cis* centromere-centromere contact maps for indicated data sets. Note the loss of the folding back in meiosis, and how the intra-arm enrichment is insulated at centromeres in meiosis.

**Extended Data Fig. 3.**
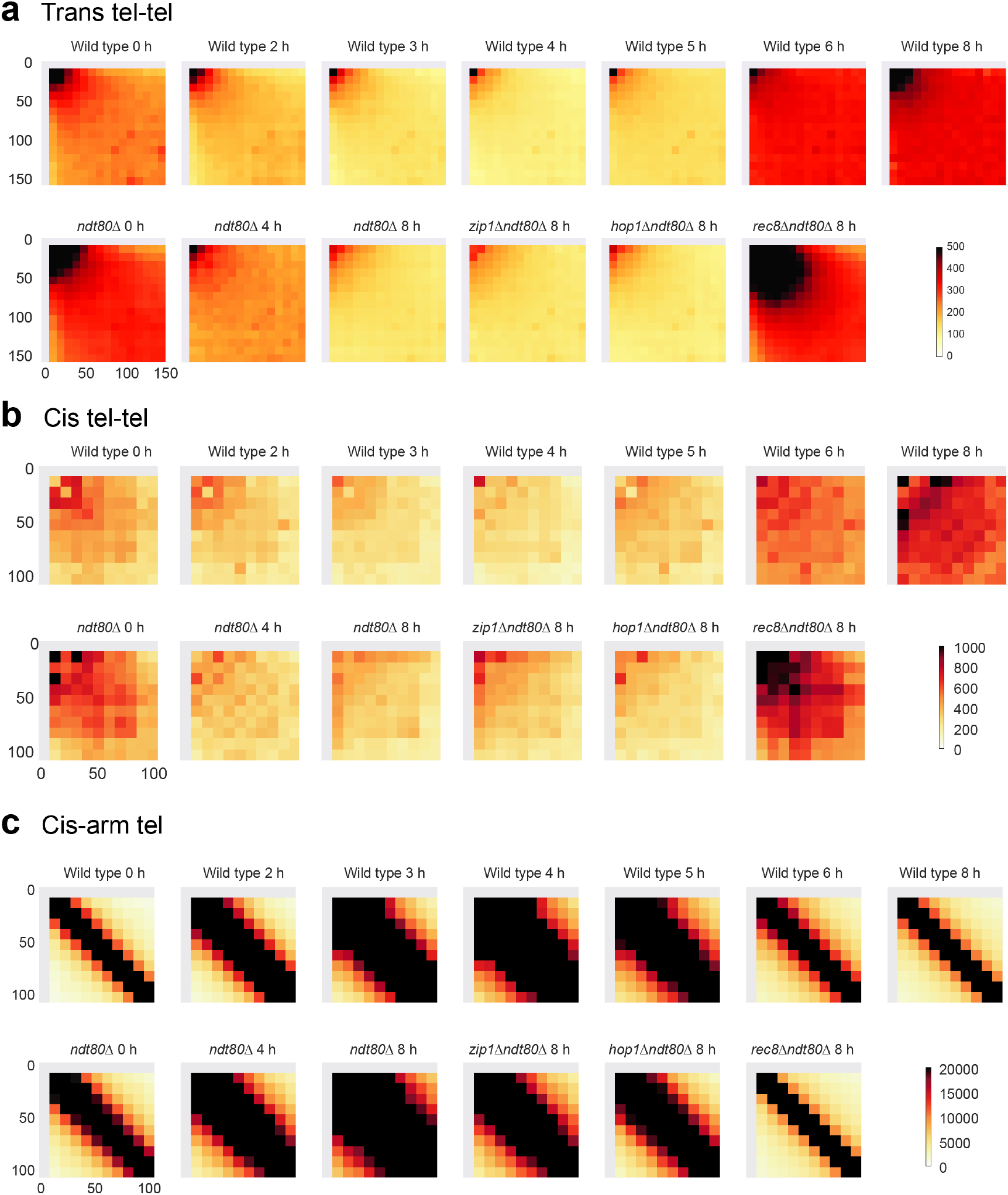
Aggregate analysis of telomeric interactions in meiosis. **a.** Average *trans* telomere-telomere contact maps for indicated datasets. **b.** Average telomere-telomere contact maps between the two telomeres of the same chromosome. **c**. Average contact map around each telomere in *cis*.

**Extended Data Fig. 4.**
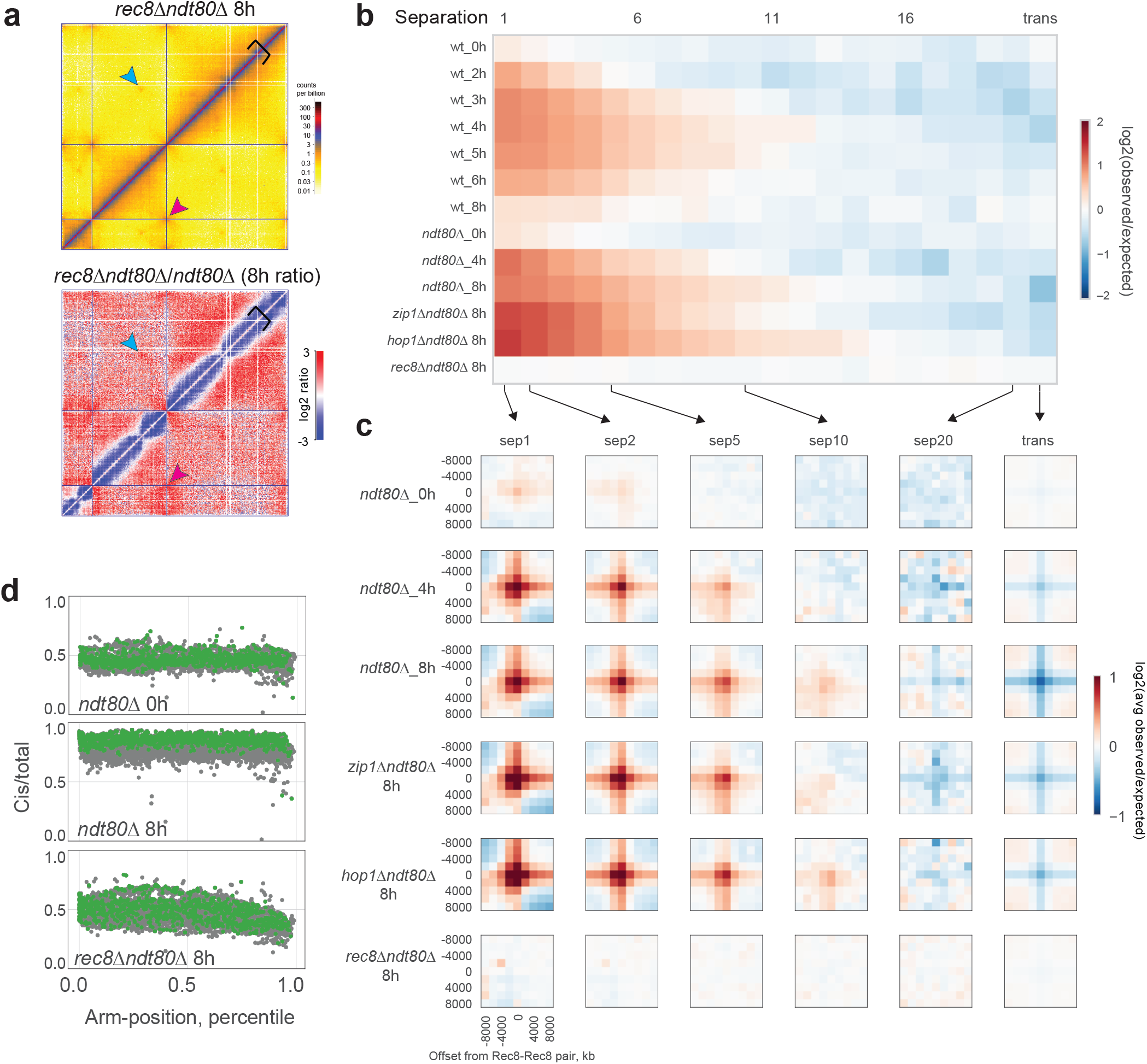
Preferred sites of Rec8 occupancy define sites of locus-specific interaction. **a.** *Left*: Hi-C contact maps of *rec8*Δ *ndt80*Δ. Chromosomes 6, 11 and 7 are shown as representatives for the whole genome. *Right*: Log2 Hi-C ratio maps of *rec8*Δ *ndt80*Δ / *ndt80*Δ. Plotted as in Fig. 1g. **b.** Log2 observed over expected contact frequency at Rec8-Rec8 peak pairs as a function of separation across datasets. **c.** Log2 observed over expected contact frequency +/-8kb around Rec8-Rec8 peak pairs at the indicated separations. Together, **b-c** demonstrate that Rec8-Rec8 enrichments are strongest between adjacent sites, decrease between non-adjacent sites with increasing genomic separation, and are absent in *trans*. Equally important, these meiotic features are lost in *rec8*Δ. As for mammalian interphase, this observation in meiosis argues for a *cis*-acting process underlying the formation of focal interactions between Rec8 sites. **d.** cis/total as a function of distance along the chromosomal arm, Rec8 sites marked in green.

**Extended Data Fig. 5.**
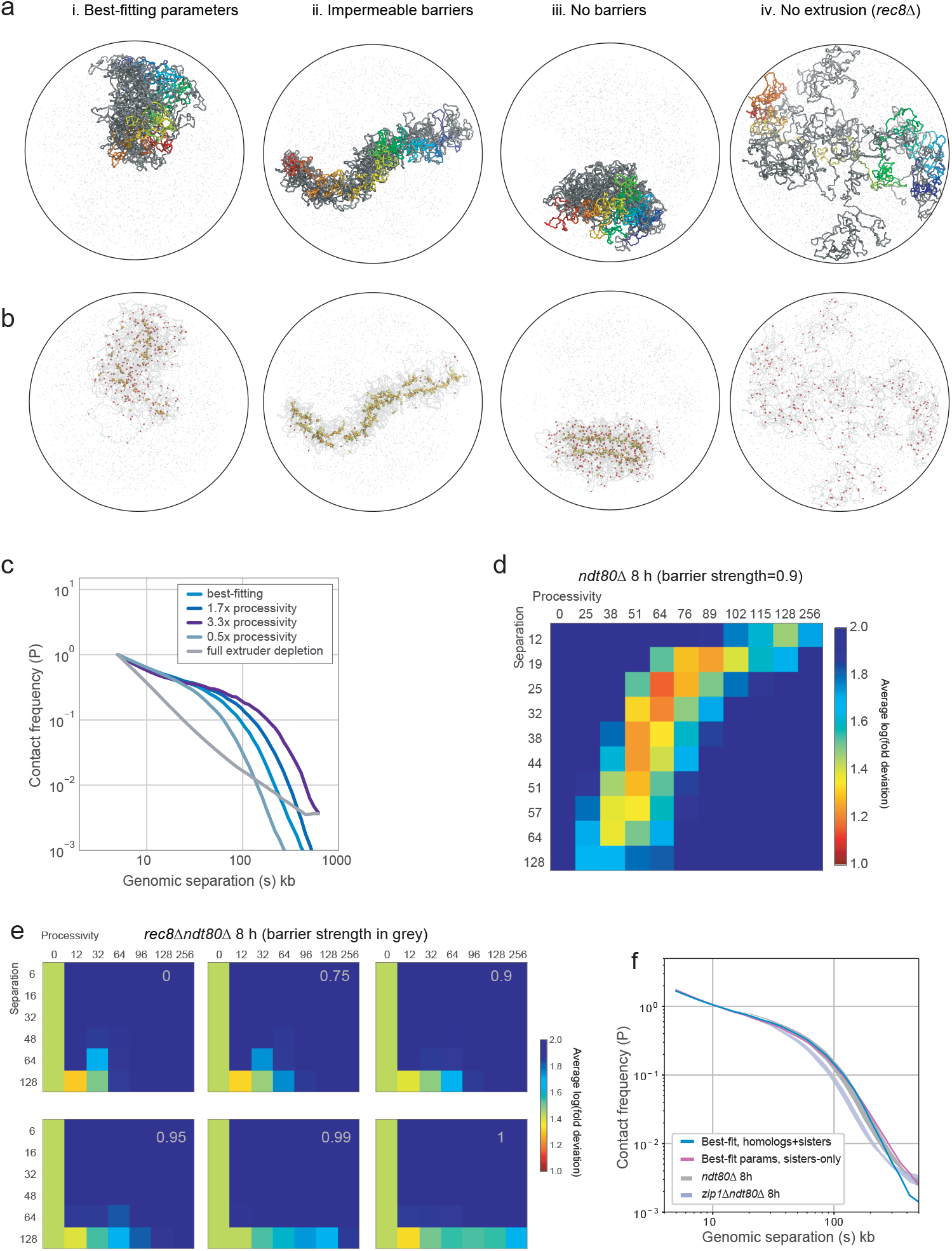
Polymer simulations of loop extrusion reveal best fitting parameters and conformations. **a.** Representative conformation for the indicated parameter sets. As in Fig. 3F, one chromatid from a homologous quartet of chromatids coloured from start to end according to the spectrum; other three coloured in grey. **b.** For the same three conformations, positions of Rec8 sites indicated with red spheres, positions of extruded loop bases in yellow, and extruders overlapping a Rec8 site in orange. Note the stable loops between neighbouring Rec8 sites creates a very elongated chromatid (ii). Also note the majority of Rec8 sites are unoccupied in (iii), despite the self-assembly of two axial cores and a strong brush. Finally, note very dispersed chromosomes in (iv), consistent with EM^3^ for *rec8*Δ. **c.** Contact frequency versus distance, *P(s)*, for indicated simulations. Note that the loss of the shoulder in *P(s)* in the case of full extruder depletion mirrors the difference between experimental *ndt80*Δ and *rec8*Δ Hi-C maps. Simulations with increased processivity predict that *P(s)* would shift rightward if unloading was impaired, as could happen in *waplΔ.* Conversely, if unloading was enhanced, simulations with decreased processivity indicate a leftward shift in P(s), until the absence of extruders. **d.** Goodness-of-fit for a fine grid of processivity versus separation at barrier strength 0.90. The best-fit occurs at similar processivity and separation as for barrier strength 0.95 shown in Fig. 3c, but with slightly lower goodness-of-fit. **e.** Goodness-of-fit to *rec8*Δ data for simulations with the indicated barrier strengths (in grey: 0.00, 0.75, 0.90, 0.95, 0.99, 1.00) over coarse grids of processivity and separation demonstrates that the best fits have few if any extruded loops, regardless of barrier strength. **f.** *P(s)* curves for simulations with sisters and homologs with the best-fitting parameters for *ndt80*Δ*-8h* maps compared to P(s) for simulations with sisters only show that simply removing homolog tethering does not recapitulate the sort of shifted P(s) seen experimentally in *zip1*Δ Hi- C.

**Extended Data Fig. 6.**
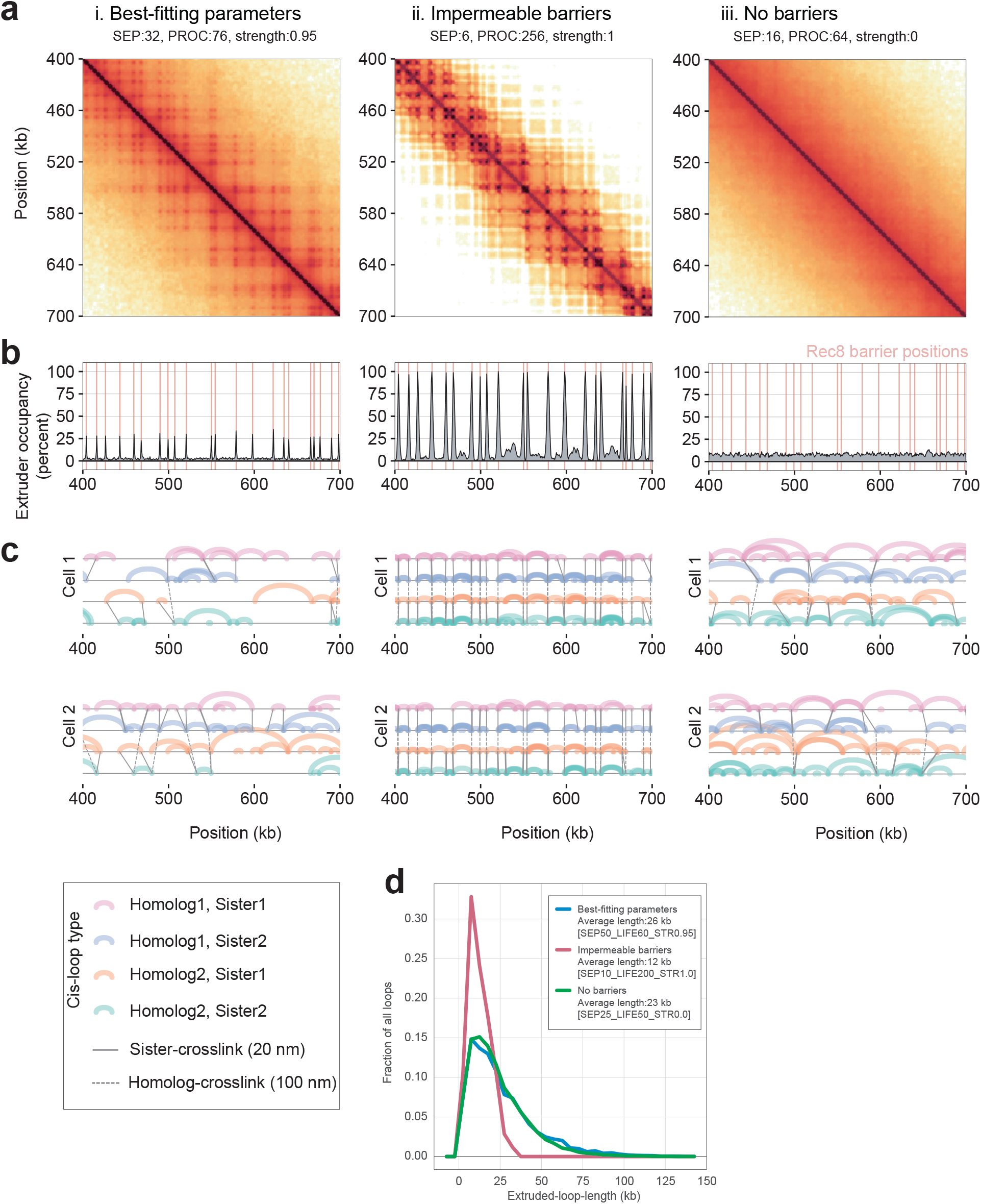
Polymer simulations of loop extrusion elucidate meiotic barrier strength. **a.** Simulated contact maps for the indicated region of chr13 for: (i) best-fitting simulations, (ii) simulations with relatively stable loops between neighboring Rec8 sites (barrier strength = 1 and high processivity), and (iii) no barriers, as in Fig. 2d**. b.** Simulated ChIP-seq profiles for the indicated region of chr13. Best-fitting simulations (i) display occupancy well below 100% at Rec8 sites. Simulations with stable loops (ii) display highly occupied Rec8 sites. Simulations without barriers (iii) have homogenous Rec8 occupancy across the genome. **c.** Positions of extruded loops (arcs) sister crosslinks (solid black lines) and homolog crosslinks (dashed lines) for four chromatids in two separate cells, showing how the simulated Hi-C maps and ChIP-seq profiles emerge from the stochastic positioning of extruded loops from cell-to-cell. For statistics, see **Supplementary Table S1**. **d.** Histogram of extruded loop lengths for indicated parameters (i, ii, iii).

**Extended Data Fig. 7.**
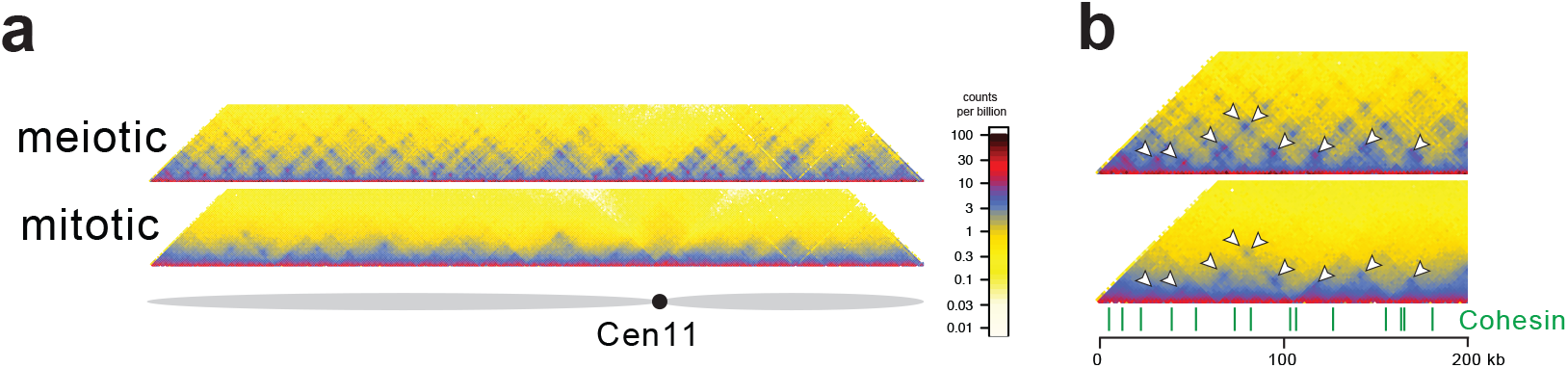
Cohesin and transcription patterns loops in meiosis and mitosis. **a.** Hi-C contact maps of chromosome 11 for meiotic (*ndt80*Δ, pachytene - top) and mitotic (wild type, nocodazole arrest - bottom) plotted at 2 kb bin resolution, showing near-diagonal interactions. Data shown is the average (n = 2). **b.** Zoom-in into contact maps on chromosome 11 (0-200kb) of *ndt80*Δ (top) and mitotic (*bottom*). Data shown is the average (n = 2). Rec8 peak sites called from ChIP-seq data^27^ are indicated in green. Arrowheads indicate sites of prominent focal interaction.

